# Polygenic risk for schizophrenia is associated with social cognition across development

**DOI:** 10.1101/034512

**Authors:** Laura Germine, Elise B. Robinson, Jordan W. Smoller, Monica E. Calkins, Tyler M. Moore, Hakon Hakonarson, Mark J. Daly, Phil H. Lee, Avram J. Holmes, Randy L. Buckner, Ruben C. Gur, Raquel E. Gur

**Affiliations:** Psychiatric and Neurodevelopmental Genetics Unit, Massachusetts General Hospital, Boston, MA 02114, USA; Stanley Center for Psychiatric Research, Broad Institute of MIT and Harvard, Cambridge, MA 02142, USA; Psychology Department, Harvard University, Cambridge, MA 02138, USA; Department of Psychiatry, Harvard Medical School, Boston, MA 02114, USA; Analytic and Translational Genetics Unit, Massachusetts General Hospital, Boston, MA 02114, USA; Medical and Population Genetics Program, Broad Institute of MIT and Harvard, Cambridge, MA 02142, USA; Neuropsychiatry Section, Perelman School of Medicine, University of Pennsylvania, Philadelphia, PA 19104, USA; Center for Applied Genomics, The Children’s Hospital of Philadelphia, Philadelphia, PA 19104, USA; Psychology Department, Yale University, New Haven, CT 06520, USA; Center for Brain Sciences, Harvard University, Cambridge, MA 02138, USA; Athinoula A. Martinos Center for Biomedical Imaging, Massachusetts General Hospital, Charlestown, MA 02129, USA

## Abstract

Breakthroughs in genomics have begun to unravel the genetic architecture of schizophrenia risk, providing methods for quantifying schizophrenia polygenic risk based on common genetic variants. Our objective in the current study was to understand the relationship between schizophrenia genetic risk variants and neurocognitive development in healthy individuals. We first used combined genomic and neurocognitive data from the Philadelphia Neurodevelopmental Cohort (PNC; 4303 participants ages 8 - 21 years) to screen 26 neurocognitive phenotypes for their association with schizophrenia polygenic risk.

Schizophrenia polygenic risk was estimated for each participant based on summary statistics from the most recent schizophrenia genome-wide association analysis (Psychiatric Genomics Consortium 2014). After correction for multiple comparisons, greater schizophrenia polygenic risk was significantly associated with reduced speed of emotion identification and verbal reasoning. These associations were significant by age 9 and there was no evidence of interaction between schizophrenia polygenic risk and age on neurocognitive performance. We then looked at the association between schizophrenia polygenic risk and emotion identification speed in the Harvard / MGH Brain Genomics Superstruct Project sample (GSP; 695 participants age 18–35 years), where we replicated the association between schizophrenia polygenic risk and emotion identification speed. These analyses provide evidence for a replicable association between polygenic risk for schizophrenia and specific aspects of neurocognitive performance. Our findings indicate that individual differences in genetic risk for schizophrenia are linked with the development of social cognition and potentially verbal reasoning, and that these associations emerge relatively early in development.

## Introduction

Schizophrenia is among the most debilitating and highly heritable of mental disorders. Recent shifts in our conceptualization of neuropsychiatric illnesses suggest that such disorders might be better understood in terms of underlying behavioral or neurobiological dimensions rather than as categories^1^. Evidence from neuroscience^2–5^, behavioral genetics^6^, and prospective clinical studies^7, 8^ suggest that schizophrenia is associated with quantitative variations in neurobiological and neurocognitive systems. These observations have led to several hypothesized relationships between schizophrenia and neurocognitive abilities^9–11^, though genetic studies examining the association between schizophrenia and IQ or educational attainment have been inconsistent ^12, 13^.

Recent genome-wide associations studies provide a window into the genetic architecture of schizophrenia, and support a complex model of psychosis liability. First, these studies demonstrate that, individually, common single nucleotide polymorphisms explain very little of the variation in schizophrenia liability^14^. The effects of common genetic variants must be taken in aggregate to explain a meaningful proportion of schizophrenia risk, indicating that the genetic liability for schizophrenia is the result of differences across many hundreds to thousands of genes and regulatory regions. GWAS data now provide the means to quantify aggregate genetic risk for schizophrenia (hereafter referred to as schizophrenia polygenic risk) for any individual, regardless of their familial or phenotypic risk^14, 15^, paving the way for a renewed interrogation of intermediate phenotypes (also known as endophenotypes)^16–18^. Understanding downstream effects of schizophrenia genetic risk variants in terms of intermediate phenotypes can highlight potential pathways that lead from genetic variation to schizophrenia, as well as potential biologically informative phenotypes to study outside of case populations^19^.

Here, we take advantage of a dataset of approximately 4300 individuals ages 8–21 collected through the Philadelphia Neurodevelopmental Cohort (PNC)^20–22^ and a replication sample of approximately 700 individuals tested as part of the Harvard / MGH Brain Genomics Superstruct Project (GSP)^23^. The PNC dataset includes genome-wide data on all participants and a comprehensive assessment of neurocognitive function across domains of general and social cognition, where measures were selected and designed to map onto specific neural circuitry.^24–26^ We took an unbiased approach to understanding the relationship between schizophrenia polygenic risk and multiple domains of neurocognitive performance, by first exploring the relationship between polygenic risk and performance across all neurocognitive measures. Genetic effects on complex traits are known to be broadly pleiotropic – a phenotypic screening approach allows us to identify the profile of associations between schizophrenia genetic risk and neurocognition, accounting for such pleiotropy within the neurocognitive phenotypes assessed. We then examined whether there was any evidence of developmentally specific effects of polygenic risk on neurocognition. Previous research has suggested that schizophrenia vulnerability might impact neurocognition at specific developmental transitions that happen during puberty and adolescence,^27, 28^ and thus associations between polygenic risk and neurocognition might be restricted to a particular age range or not begin until a critical developmental period begins. Using the resources of the PNC and GSP, we explore and replicate tests of the hypothesis that individual differences in schizophrenia genetic risk are related to quantitative dimensions of neurocognition.

## Methods

### Participants

Our primary analytic sample was drawn from the Philadelphia Neurodevelopmental Cohort (PNC), a Children’s Hospital of Philadelphia health network-based sample of about 9500 individuals aged 8–21 years from the greater Philadelphia area (details in Calkins et al. and Gur et al.)^20, 24^. Participants who provided assent / consent gave genetic samples and written permission to be recontacted for further research. The University of Pennsylvania and Children’s Hospital of Philadelphia Institutional Review Boards approved the study. After stratification based on sex, age, and ethnicity, PNC participants were recruited through random selection. Inclusion criteria were (1) ability to provide informed consent (parental consent where age < 18), (2) English proficiency, and (3) physical and cognitive functioning sufficient to complete clinical assessment interviews and cognitive testing on a computer.

Our replication sample was drawn from the Harvard / MGH Brain Genomics Superstruct Project (GSP). The GSP is a study cohort that includes neuroimaging, genomic, and cognitive data on over 4000 healthy participants^23^. The present sample included an age-restricted subsample that performed behavioral tasks compatible with the discovered effects in the PNC. Because of differences in protocols, only a subset of PNC assessments could be tested for replication. All GSP participants provided written informed consent for biomedical research approved by the Partners Healthcare Institutional Review Board or the Harvard University Committee on the Use of Human Subjects. Inclusion criteria were (1) English proficiency, (2) age 18–35 years, (3) no history of psychiatric illness or major health problems, and (4) physical and cognitive functioning sufficient to complete MRI scanning and cognitive testing.

### Genetic analysis

As polygenic risk scores are sensitive to ancestry, we restricted our analysis to genotypes individuals with self-described White non-Hispanic ancestry. Data cleaning and imputation were performed using standard procedures (see Robinson et al.^21^ for PNC data and Hibar et al.^29^ for GSP data). In the PNC dataset, the ancestry threshold was relaxed to a pi_hat of 0.1 (0.125 for the GSP dataset), which is a level of relatedness equal to or less than that of first cousins. Imputed data were used to generate individual schizophrenia polygenic risk scores using the procedures described in Purcell et al.^15^ and Ripke et al.^14^. In brief, polygenic risk scores estimate genome-wide common variant liability for a trait through a weighted sum of many thousands of risk alleles. The polygenic risk score used here was generated using summary statistics from the Psychiatric Genomics Consortium (PGC) recent meta-analysis of schizophrenia^14^, and includes only SNPs with p < 0.05. This version of the score was selected because it most commonly maximized the schizophrenia risk explained in independent case-control samples (approximately 20% of case-control variation)^14^. In addition to limiting the analysis to participants of European descent, the first ten principal components of ancestry were controlled for in all analyses.

### Phenotypic Neurocognitive Assessment and Analysis

PNC participants completed the Computerized Neurocognitive Battery (CNB)^24, 25^. The CNB was developed from tasks that map onto specific brain systems, as identified through functional neuroimaging^26^. Psychometric properties and task descriptions for the CNB measures are included elsewhere^21, 24–26, 30^. The CNB provides accuracy and speed measures of: executive function (abstraction and mental flexibility, attention, working memory), memory (verbal, spatial, facial), complex cognition (verbal reasoning, nonverbal reasoning, spatial processing), social cognition (emotion identification, emotion differentiation, age differentiation) as well as speed measures for sensorimotor and pure motor function.

GSP participants completed an online battery of personality, cognitive, and behavioral measures assessing a broad range of domains. A full list of measures and details of neurocognitive assessment for the GSP are described in Holmes et al. ^23^. A subset of measures that are available in the PNC were also available in the GSP dataset (in comparable or identical form).

Differences in performance attributable to age and sex were regressed out of all neurocognitive variables (speed and accuracy) prior to analysis. Any individual scores more than four standard deviations from the mean for a particular test were excluded. Linear regression was then used to examine the relationship between schizophrenia polygenic risk and neurocognitive performance. In our primary analytic sample (PNC), Bonferroni correction was applied to correct for the number of comparisons across all neurocognitive variables (26 comparisons: 12 accuracy variables and 14 speed variables) with an alpha threshold of 0.05. This correction is conservative given the correlation structure of the neurocognitive phenotypes^21^.

## Results

The final analytic sample from the PNC included 4303 participants (50% female) ranging in age from 8–21 (near uniform distribution with mean age of 13.8 years). As described in Robinson et al. ^21^, most of the 838 genotypes, white non-hispanic PNC individuals excluded from this analysis were removed for outlying ethnicity or relatedness to another person in the dataset. After Bonferroni correction, two neurocognitive measures were significantly associated with schizophrenia polygenic risk at p < 0.05 (see Figure 1a). These were verbal reasoning speed (analogies) (β = −0.058, p < 5E-4 uncorrected) and emotion identification speed (β = − 0.066, p < 5E-5 uncorrected). For both variables, increases in polygenic risk were related to linear decreases in speed of responses, with the highest quartile showing the slowest speeds (longest reaction times) and the lowest quartile showing the highest speeds (shortest reaction time) (see Figure 1b). This was not the case for other variables related to response speed, for example sensorimotor speed, where no differences were observed between the lowest and highest quartiles of schizophrenia genetic risk (see Figure 1b). Schizophrenia polygenic risk was unrelated to matrix (nonverbal) reasoning ability, a common proxy for IQ (no relationship after correction for multiple comparisons; p = 0.035 uncorrected) or general cognitive ability (i.e. general intelligence or ‘g’, based on factor analysis; p= 0.64 uncorrected).

**Figure 1.**
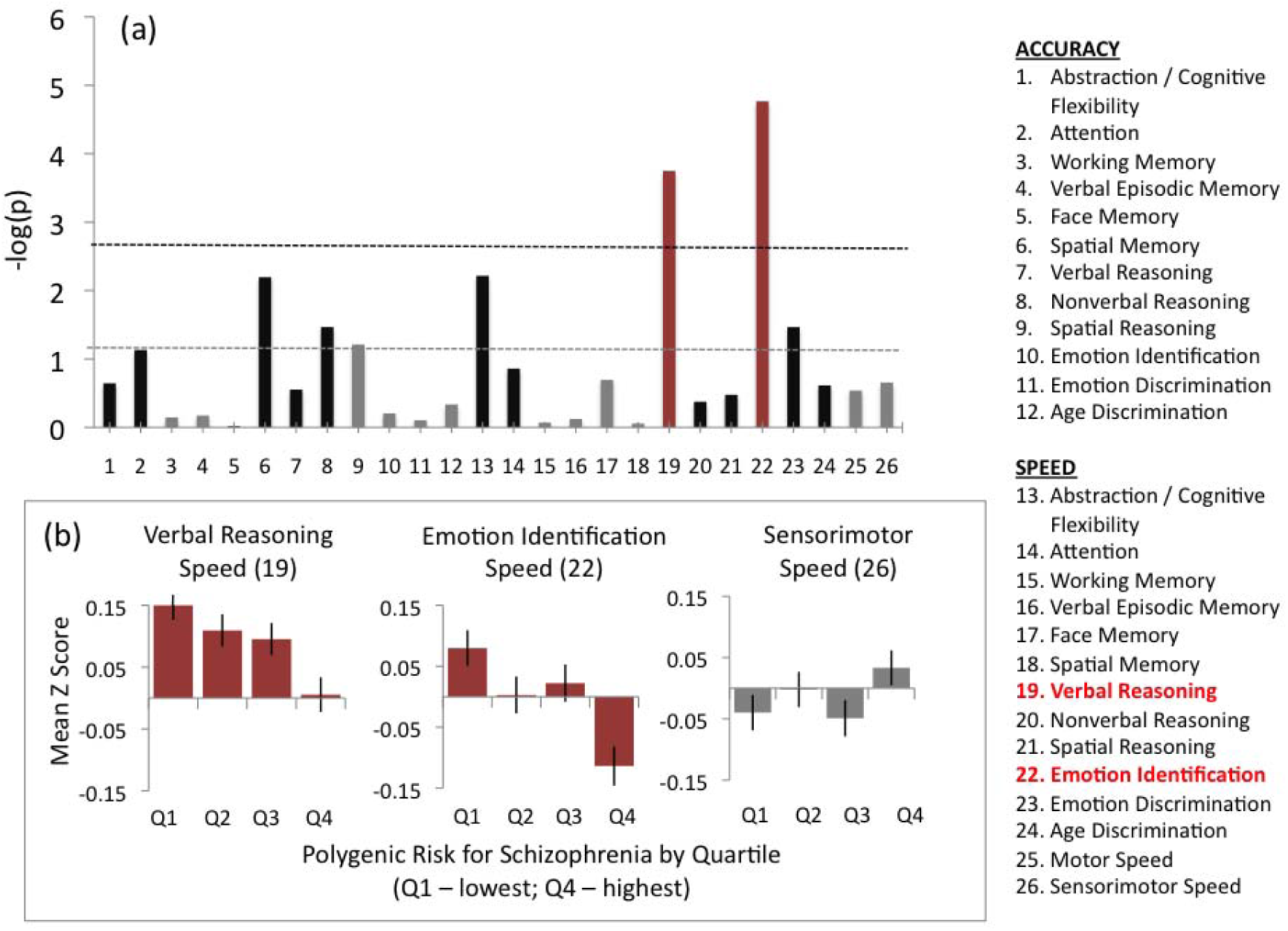
Schizophrenia polygene scores and neurocognitive performance. (a) Linear regression was used to estimate associations between schizophrenia polygenic risk, estimated from genome-wide data, and performance for each neurocognitive variable (labels shown on the right). To best illustrate the strength of the evidence for each association, relationships are plotted in terms of the negative base-10 logarithm of the p-value, when regressing neurocognitive performance on schizophrenia polygenic risk estimates for each participant. The gray line shows the threshold for statistical significance based on p < 0.05, uncorrected. The black line shows the threshold for statistical significance after Bonferroni correction for all 26 comparisons (p < 0.05 corrected). Red bars show variables where an association exceeded the threshold for statistical significance (verbal reasoning speed and emotion identification speed). For nonsignificant associations, black bars indicate a negative relationship between schizophrenia polygenic risk and neurocognitive performance (i.e. greater polygenic risk associated with poorer performance) and gray bars indicate a positive relationship. (b) The relationship between schizophrenia polygenic risk and neurocognitive performance is shown for the two speed variables (emotion identification and verbal reasoning) where associations were statistically significant after correction for multiple statistical tests and (for comparison purposes) associations with a general measure of response speed. Participants from the primary analytic sample (PNC dataset) were divided into four groups of approximately equal size based on level of schizophrenia polygenic risk. Quartile 1 (Q1) includes individuals with the lowest schizophrenia polygenic risk. Quartile 4 (Q4) includes individuals with the highest schizophrenia polygenic risk. Mean z-score is plotted on the y-axis, with higher values reflecting better performance. For both emotion identification and verbal reasoning speed, increasing polygenic risk was linearly associated with decrease in neurocognitive performance.

To understand specificity of the relationship between schizophrenia polygenic risk and speed of verbal reasoning and emotion identification, we performed the same analysis, controlling for overall motor speed, sensorimotor speed, as well as matrix/nonverbal reasoning ability. Both associations were similar after controlling for motor speed and sensorimotor speed (emotion identification speed: β = −0.079, p < 5E-7 uncorrected; verbal reasoning speed: β = −0.065, p < 5E-5 uncorrected) and matrix reasoning ability (emotion identification speed: β = −0.065, p < 5E-5 uncorrected; verbal reasoning speed: β = −0.057, p < 5E-4 uncorrected).

To understand the impact of speed accuracy trade-offs on the strength of each association, we also looked at both emotion identification speed and verbal reasoning speed, controlling for emotion identification accuracy and verbal reasoning accuracy, respectively. Emotion identification speed was associated with schizophrenia polygenic risk, even after controlling for emotion identification accuracy (β = −0.065, p < 5E-5 uncorrected). Verbal reasoning speed was also associated with schizophrenia polygenic risk after controlling for verbal reasoning accuracy (β = −0.056, p < 5E-4 uncorrected).

For both variables, schizophrenia polygenic risk accounted for approximately 0.3 – 0.5% of the variation in neurocognitive performance.

### Developmental Specificity

We evaluated the relationship between polygenic risk and speed at each year of age (see Figure 2). There was no evidence that the associations between schizophrenia polygenic risk and emotion identification, or between schizophrenia polygenic risk and verbal reasoning, were emerging later in development (e.g. after puberty). There was also no evidence for the opposite pattern – early associations that disappeared later in development due to potential compensatory changes. The relationship between schizophrenia polygenic risk and neurocognitive performance was statistically significant by age 8 for verbal reasoning speed (β = −0.094, p < 5E-2) and by age 9 for emotion identification speed (β = −0.11, p < 5E-2). In both cases, there was no interaction between schizophrenia polygenic risk and age for either emotion identification speed (p = 0.38) or verbal reasoning speed (p = 0.11). Thus we find no evidence for modulation in the relationship between schizophrenia polygenic risk and emotion identification or schizophrenia polygenic risk and verbal reasoning across development.

**Figure 2.**
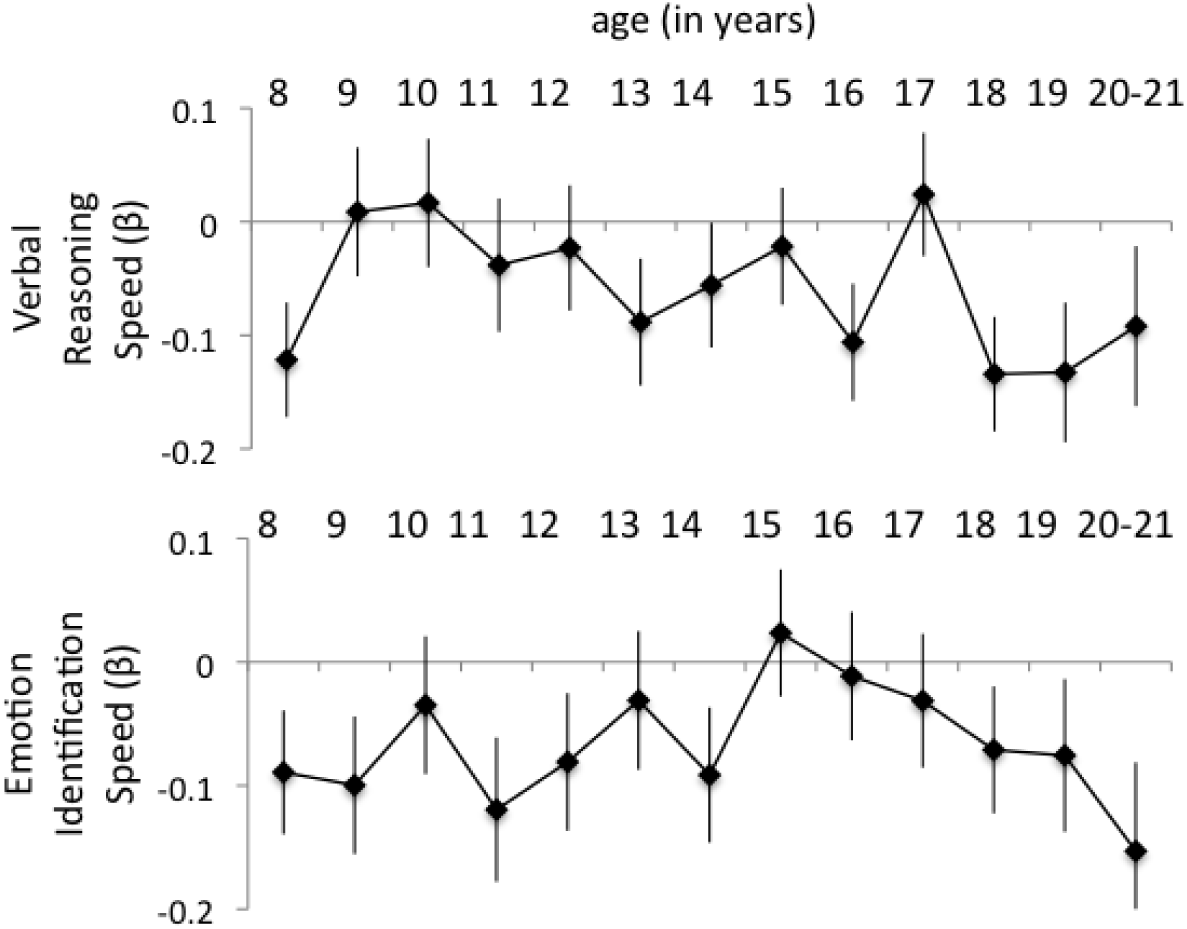
Schizophrenia polygene scores as predictors of emotion identification speed and verbal reasoning speed, across age. Shown are associations between schizophrenia polygenic risk and speed of emotion identification and verbal reasoning, at each year of age. Bars give +/− 1 standard error of the effect size estimate. Although both measures were significantly associated with schizophrenia polygenic risk, there was no significant interaction of polygenic risk and age on neurocognitive performance for either variable.

### Replication in an Independent Sample

In the GSP dataset, there were 695 participants completed the emotion identification measures used in the CNB and met our inclusion criteria. These participants were 53% female and ranged in age from 18–35 (mean: 21.5; SD: 3.2). This independent sample allowed us to test replication of the relationship between schizophrenia polygenic risk and emotion identification speed. The stimuli for the emotion identification task were shared from the CNB at the initiation of the GSP data collection effort, allowing for a true, independent replication using the same stimuli and test in an independent cohort. The association between schizophrenia polygenic risk and emotion identification speed was replicated, despite GSP participants being sampled from a very different population (β = −0.09, p < 0.05). This association again survived controlling for IQ (as estimated by matrix reasoning performance) and a measure of psychomotor response speed. Verbal reasoning speed was not one of the neurocognitive measures included in the GSP, so we were unable to assess replication for this association.

**Figure 3.**
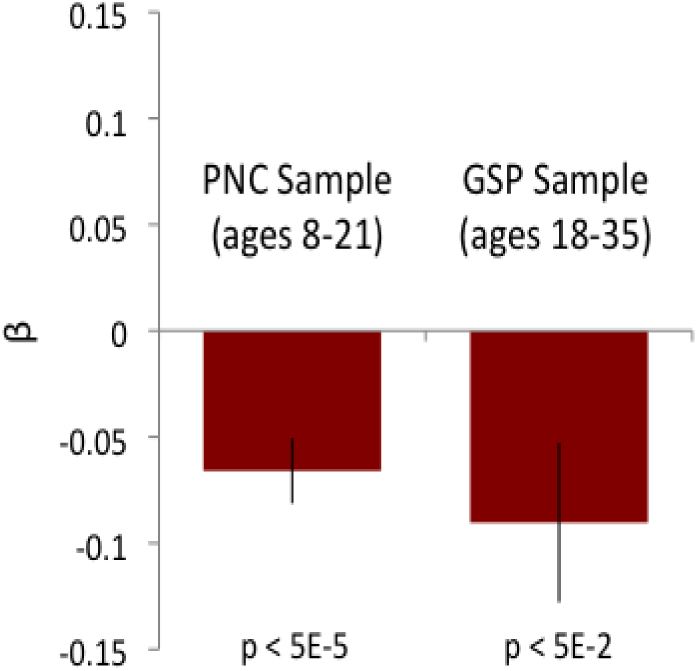
Schizophrenia polygenic risk and emotion identification speed: Replication in an independent sample of adults. Shown are effect size relationships between emotion identification speed (controlling for age and sex) and schizophrenia polygenic risk in the original discovery sample (PNC ages 8–21) and the replication sample (GSP ages 18–35).

## Discussion

In two large samples, spanning middle childhood to adulthood, we identify a small, but statistically replicable relationship between schizophrenia polygenic risk and speed of emotion identification. We also found an equally significant association between schizophrenia polygenic risk and speed of verbal reasoning in our developmental sample (ages 8 – 21). Both associations survived correction for multiple comparisons in our developmental (discovery) sample and the inclusion of covariates related to general intelligence and sensorimotor speed. Critically, the association between schizophrenia risk and emotion identification replicated in an independent, demographically distinct sample of adults.

We found no evidence that either association depended on developmental phase: effects were significant by age 9 and effect sizes were consistent across middle childhood, adolescence, and early adulthood. These results suggest a relatively early perturbation affecting neurocognitive development that is present before the typical onset of psychotic illness in young adulthood. Finally, we found that even intermediate levels of schizophrenia polygenic risk were associated with reductions in emotion identification and verbal reasoning speed. In other words, the relationship between schizophrenia polygenic risk and performance was evident across the spectrum of polygenic risk, as opposed to being observed only in those at the highest polygenic risk^31^. Our results indicate that common variants that increase risk for schizophrenia impact the development of social cognition and possibly verbal reasoning.

The finding of a relationship between schizophrenia polygenic risk and emotion identification performance is consistent with a large body of literature emphasizing the importance of social cognition in schizophrenia, individuals who go on to develop schizophrenia, and individuals at risk of schizophrenia. Schizophrenia is associated with profound and consistent deficits in social cognition^32–35^. These deficits appear early, sometimes decades before the onset of illness^36–38^ and are strongly related to core aspects of symptomatology and everyday social functioning ^39–43^. Differences in social cognition are observable in healthy individuals with a potential genetic predisposition towards developing schizophrenia ^44–46^. Early environmental factors linked with the development of schizophrenia ^47, 48^ are also associated with adult differences in social cognition^49^. Even in healthy populations, quantitative differences in psychosis-like characteristics are linearly related to differences in emotion identification^31^. There is extensive literature with functional neuroimaging indicating abnormalities in regional brain activation to emotion identification tasks in patients with schizophrenia ^4, 5, 50^ and those with psychosis spectrum features^51^, including individuals from the PNC who underwent neuroimaging^52^. Our specific findings provide further support, based on polygenic risk estimates, that schizophrenia is a disorder that is fundamentally related to the development of social cognition.

Abnormalities in verbal reasoning have also been identified in schizophrenia and first-degree relatives of schizophrenia patients^53^. Although this association was not available for replication due to the absence of a comparable phenotype in our replication sample, the effect size difference was comparable to our finding relating polygenic risk and emotion identification speed. Perhaps deficits in processing emotion and verbal communication combine in creating difficulties in social communication, contributing to social functioning deficits related to schizophrenia risk. Abnormalities in the development of social understanding and communication – in the presence of other life events or cognitive vulnerabilities – could set the stage for psychosis later in life. This possibility is consistent with findings in the entire PNC sample that individuals with psychosis spectrum features are delayed in their neurocognitive development already by age 8^54^. Notably, the most pronounced delays were in complex reasoning and social cognition.

These results have implications for our understanding of schizophrenia genetic risk in relation to neurocognition, particularly social cognition. First, our findings validate the notion that risk genes might perturb cognition in a dimensional fashion -- with a linear (as opposed to threshold) relationship between genetic load and cognitive impairment. We found such a relationship for both emotion identification speed and verbal reasoning speed, indicating that quantitative dimensions of genetic risk and cognitive vulnerability can provide critical information for understanding psychopathology. Further, that these relationships did emerge so early may make it unlikely that social deficits arise as a consequence of the expression of psychotic or psychosis-like characteristics (e.g. stigma or social rejection could drive social isolation and subsequent deterioration of social cognitive skills in patients with schizophrenia). Indeed, the differences at even low levels of polygenic risk point to a more direct and fundamental relationship.

It is noteworthy that our findings were mainly for speed of processing rather than accuracy. This effect is consistent with earlier studies that showed reduced speed in individuals genetically related to probands with schizophrenia and may indicate compensatory strategies to mitigate vulnerability^55^. Furthermore, processing speed has been implicated in meta-analyses as a major deficit domain in schizophrenia^11^. Longitudinal studies are needed to examine whether reduced speed of processing during development portends deficits that extend to accuracy and eventually to schizophrenia.

The present study has several limitations. The wide age range spanning from childhood to early adulthood is an epoch associated with rapid improvement in cognitive performance. While enabling the identification of developmental effects, this characteristic of the PNC may mask effects that could be detected in equally sized samples of adults with a narrower age range. The effects we report, while significant, are small – accounting for only between 0.5 – 1% of variance in neurocognitive performance. Given the small effect size, the association between schizophrenia polygenic risk and neurocognitive performance reported is primarily of theoretical interest by illustrating how polygenic risk is linked to a specific dimensional cognitive domain. The small effects suggests that the link as measured will not be useful for prediction or making conclusions about individuals. The size of the identified associations may also reflect the overall normative nature of the sample; estimated risk could still confer clinically significant effects in vulnerable individuals. Finally, only one of the effects could be tested in an independent sample because the other measure was not available, underscoring the need for harmonizing measures across genomic projects.

## Conclusion

Notwithstanding these limitations, our results highlight a new and important role for intermediate phenotypes in the GWAS-era: as a means of understanding mechanism from validated genetic predictors of disease^19^. In this role, intermediate phenotypes are not only useful but also fundamental, as they allow us to understand how genes contribute to the development of disease. Finally, our results point to one potential mechanism linking genetic risk for schizophrenia to psychosis through alterations in social abilities. These alterations arise early in development before typical onset of psychosis. This finding is consistent with decades of research on the fundamental nature of social deficits in schizophrenia, further indicating that these social deficits may arise from schizophrenia risk-related common genetic variants. The finding that schizophrenia has a genetic basis in individual differences opens up many pathways for further study, including the translation of knowledge from social neuroscience and social cognitive psychology towards the elucidation of the roots of major psychiatric illness.

## Acknowledgements

This study was supported by National Institutes of Health grants F32MH102971 (Dr Germine), MH089983 and MH096891 (Dr. Gur) and MH089924 (Dr. Hakonarson), K08MH079364 (Dr. Calkins), K99MH101367 (Dr. Lee), U01MH094432 (Dr. Daly), and K01MH099286 (Dr. Robinson),

